# Salt-dependent self-association of trinucleotide repeat RNA sequences

**DOI:** 10.1101/2023.11.26.568751

**Authors:** Hiranmay Maity, Hung T. Nguyen, Naoto Hori, D. Thirumalai

**Affiliations:** Department of Chemistry, University of Texas at Austin, USA; Department of Chemistry, University at Buffalo, NY, USA; School of Pharmacy, University of Nottingham, Nottingham, NG72RD, United Kingdom; Department of Physics, University of Texas at Austin, USA

## Abstract

Low complexity repeat RNA sequences self-associate by homotypic interactions to form condensates. Using simulations of a coarse grained Single-Interaction Site model for (CAG)_n_ (*n* = 30 and 31), we show that the salt-dependent free energy gap, ∆*G*_*S*_, between the ground (perfect hairpin) and the excited state (slipped hairpin (SH) with one CAG overhang) of monomer (*n* even) is the primary factor that determines the rates and yield of self-assembly. For odd *n*, the SH ground state free energy (*G*_*S*_) is used to predict self-association kinetics. As the monovalent salt concentration, *C*_*S*_, increases ∆*G*_*S*_ and *G*_*S*_ increases, which in turn decreases the self-association rates. In contrast, ∆*G*_*S*_ for scrambled sequences, with the same length and sequence composition as (CAG)_31_ but with higher complexity, is larger which greatly suppresses the propensities to aggregate. Although demonstrated explicitly for (CAG)_30_ and (CAG)_31_ polymers, the finding that there is an inverse correlation between *C*_*S*_-dependent ∆*G*_*S*_ or *G*_*S*_ and RNA aggregation is general. Our predictions are amenable to experimental tests.

## Introduction

A series of experiments^1–8^ have established that low complexity repeat RNA sequences, such as (CAG)_n_ and (CUG)_n_, (n is the number of repeat units) undergo phase separation. In the two phase region, the high density droplet coexists with the sol or low density dispersed phase. The qualitative features of the phase separation may be understood using the venerable Flory-Huggins theory, ^9,10^ although in RNA temperature is not as relevant as salt concentration.

Besides their intrinsic biological interest, the transcribed products of (CAG)_n_ and (CUG)_n_ are implicated in Huntington’s disease, muscular dystrophy and amytropic lateral sclerosis.^11–13^ Recently, using coarse-grained simulations and theoretical arguments, we provided a conceptual framework for describing condensate formation in repeat nucleotide sequences.^14,15^ The driving force for self association arises both from favorable intermolecular Watson-Crick (WC) base pair formation as well as the degeneracy associated with a large number of ways such base pairs can form^14,16^ in a droplet. In a recent account,^15^ we showed that the propensity of repeat RNA polymers to aggregate can be inferred from the free energy spectrum of the monomer. For even *n* the ground state of (CAG)_n_ polymer is a perfect hairpin (PH) that is stabilized by base stacking and Watson-Crick base pair formation. In this case, we showed that the propensity for self-association between (CAG)_n_ polymer is determined by the free energy gap, ∆*G*_*S*_, between the ground state (GS) and the excited state in which at least one (CAG) unit is exposed, resulting in slipped hairpin (SH) states. The self-association propensity decreases as ∆*G*_*S*_ increases. For odd *n*, the GS is already in the SH state, which enhances the propensity to self-associate. Our theoretical prediction that (CAG)_2n+1_ should have higher tendency to aggregate than (CAG)_2n_ was validated using computer simulations at 0.1 M monovalent salt concentration. ^15^ A recent all atom molecular dynamics simulations of homopolymeric RNA sequences^17^ have also suggested that single chain properties could reveal the propensity to undergo phase separation.

The spectrum of states sampled by the (CAG)_n_ polymers may be altered by varying the external conditions. Here, we explored the extent to which aggregation of (CAG)_n_ changes as the salt concentration is varied. We simulated the two sequences (CAG)_30_ and (CAG)_31_ both of which undergo phase separation at high densities. The population of aggregation prone hairpin state of the RNAs is suppressed as the salt concentration, *C*_*S*_, increases from 0.15 M to 0.5 M. The rate of formation of dimer decreases with an increase in *C*_*S*_, which is in accord with the theory that ∆*G*_*S*_ increases with increasing *C*_*S*_. In contrast, the propensity to self-associate decreases in scrambled sequences with identical composition because ∆*G*_*S*_ increases substantially.

## Results and Discussion

### Salt modulates the population of different hairpin structures

We performed low friction Langevin dynamics simulations (see Supporting Information (SI)) at 37^*◦*^ C by varying the salt concentration, *C*_*S*_, from 0.15 M to 0.5 M. The secondary structures obtained at these conditions are classified broadly into stem-loop or hairpin-like structures. The terminal nucleotides in the hairpin-like structures are spatially close. Single molecule Foster Resonance Energy Transfer (smFRET) spectroscopy characterized the structures of trinucleotide repeat DNA hairpins ((CTG)_*n*_ ^18^ and (CAG)_*n*_ ^19^). It was found that the most populated state for (CAG)_14_ (the 5^*′*^ and 3^*′*^ ends are close) is a perfect hairpin whereas the ground state of (CAG)_15_ is slipped (FRET efficiency is lower compared to the ground state of (CAG)_14_, thus exposing one CAG unit).

Because the FRET efficiency is related to the distribution of the end-to-end distance, we calculated *P* (*R*_*ee*_) for both A(CAG)_30_A and A(CAG)_31_A at *C*_*S*_ = 0.15 M and 0.5 M (Figures 1A and 1B). We find that *P* (*R*_*ee*_) has more than one peak indicating that multiple stem-loop conformations are sampled. The peak at *R*_*ee*_ = 1.45 nm is approximately at the equilibrium base pair distance (≈ 1.38 nm) between two beads, showing that the terminal nucleotides are in spatial proximity. The peaks located at *R*_*ee*_ *>* 1.45 nm are either due to break down of terminal G-C base pair (bp) or slippage in the strands. As the *C*_*S*_ value is increased to 0.5 M from 0.15 M, the amplitude of the first peak in *P* (*R*_*ee*_) increases for both the sequences, which implies that increment in salt concentration stabilizes the base pair strongly.

**Figure 1:**
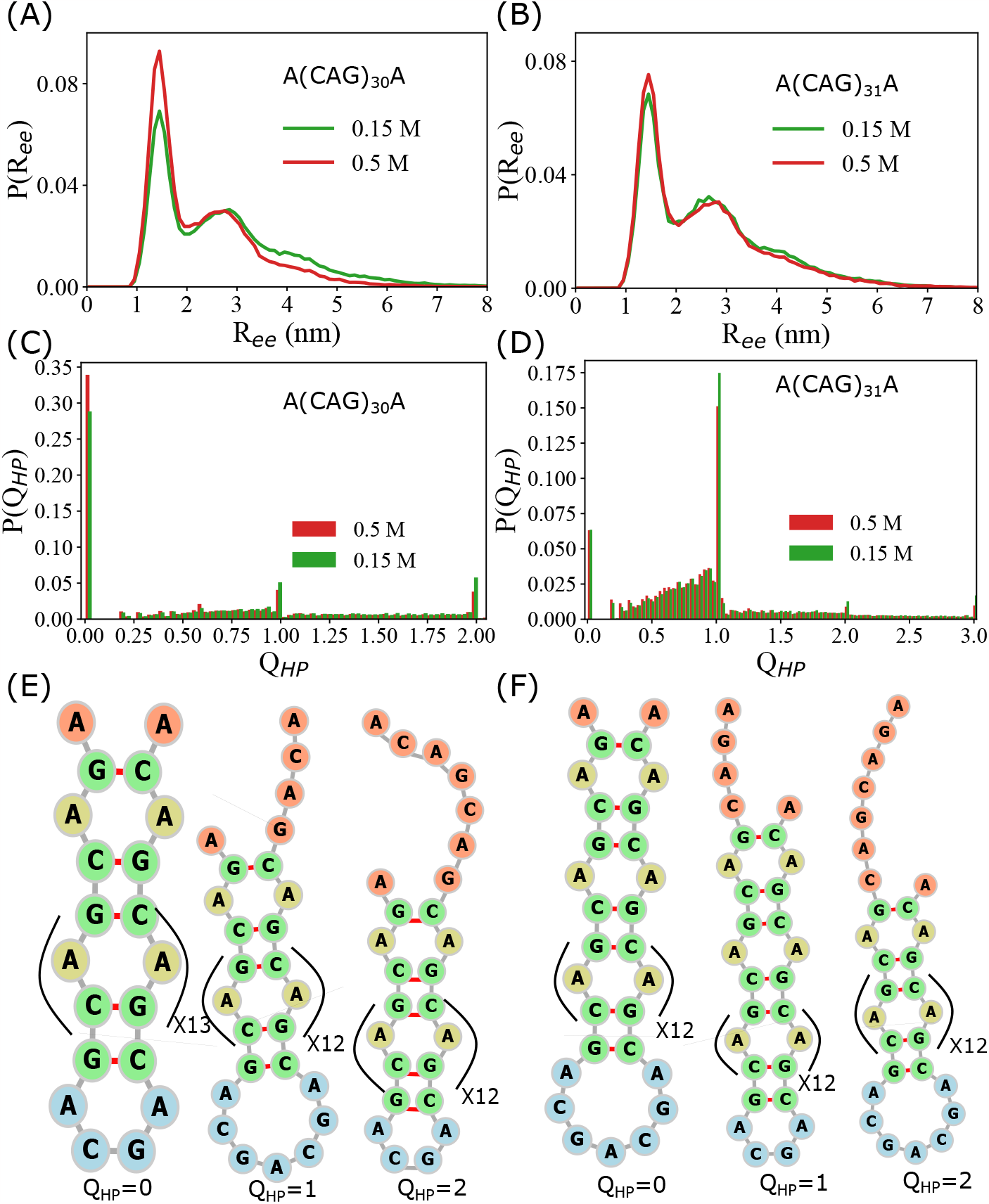
Characterization of hairpin structures: (A) Distribution, *P* (*R*_*ee*_), of the end-to-end distance, *R*_*ee*_, for A(CAG)_30_A at salt concentrations, *C*_*S*_ = 0.15 M and 0.5 M are in green and red, respectively. (B) *P* (*R*_*ee*_) as a function of *R*_*ee*_ for A(CAG)_31_A at *C*_*S*_ = 0.15 M and 0.5 M are in green and red, respectively. Multiple peaks in the *P* (*R*_*ee*_) are signatures of distinct stem-loop conformations in the ensemble of hairpin structures. (C) The probability distribution *P* (*Q*_*HP*_) of *Q*_*HP*_ for (CAG)_30_ at *C*_*S*_ = 0.5 M and 0.15 M are in red and green, respectively. As *C*_*S*_ increases the probability of *Q*_*HP*_ = 0 increases whereas *P* (*Q*_*HP*_) with *Q*_*HP*_ *>* 0 decreases. (D) *P* (*Q*_*HP*_) for (CAG)_31_ at *C*_*S*_ = 0.15 M (green), 0.5 M (red) shows that the ground state population of (CAG)_31_ decreases, i.e. *P* (*Q*_*HP*_) with *Q*_*HP*_ = 1 decreases. (E) Hairpin structures with *Q*_*HP*_ = 0, 1, and 2 for (CAG)_30_. (F) Same as (E) except the results are for (CAG)_31_. The hairpin structures are generated using forna^32^

In order to elucidate the microscopic structures of the stem-loop conformations, we computed the order parameter, *Q*_*HP*_, for a given conformation using the Eq. 2 by accounting for the arrangements of the bps in the stem region. *Q*_*HP*_ measures the deviation in the arrangement of base pairs in the stem region relative to the the arrangement in the perfect hairpin (PH) structure in which there is no mismatches, except for the unavoidable A-A mismatches. An ensemble of structures with *Q*_*HP*_ = 0 is, therefore, identical to the PH (Figure 1E and 1F). The set of bps, *S*_*bp*_, representing the stem of a PH can be expressed as, *S*_*bp*_ = (*i, j*) : *i* + *j* = *N*_*T*_ + 1, where, *i* and *j* are the indices of the nucleotides forming the base pair, and *N*_*T*_ is the number of nucleotides in the sequence. Hairpins with *Q*_*HP*_ = *m*, where, *m* is a positive integer, signify the strand slippage by *m* repeat units of CAG from either the 5^*′*^ or 3^*′*^ end. Fractional values of *Q*_*HP*_ indicate the formation of one or more than one bulge at the stem region of the hairpins (Figure S3).

To investigate the effects of salt on the population of different hairpin states, we calculated the distribution, *P* (*Q*_*HP*_), of *Q*_*HP*_ at *C*_*S*_ ranging between 0.15 and 0.5 M (Figure 1C, 1D and S1 in SI). The most populated structures or the ground state (GS) conformations of (CAG)_30_ is a PH, whereas it is a slipped hairpin (SH) with one unit of CAG overhang at the terminal for (CAG)_31_. Alternation in GS conformation between PH and SH in going from an even to an odd repeats has been observed in both repeat DNA sequences in experiments^19,20^ and in simulations^15^ of repeat RNA sequences. Although, the GS remains PH (SH) for (CAG)_30_ ((CAG)_31_) at all *C*_*S*_ values, its population changes as *C*_*S*_ value is varied. For instance, the population of the PH of (CAG)_30_ changes to ≈ 0.33 from ≈ 0.28 as *C*_*S*_ is increased to 0.5 M from 0.15 M. Notably, there is a decrease in *P* (*Q*_*HP*_ = 2) and *P* (*Q*_*HP*_ = 1), which shows that the population of slipped hairpin decreases as *C*_*S*_ increases. Interestingly, we also found a suppression in the population in *P* (*Q*_*HP*_ = 1) for (CAG)_31_. Combining the results for (CAG)_30_ and (CAG)_31_, we conclude that slippage in strands is reduced as *C*_*S*_ increases. Experiments probing the slippage dynamics of (CAG)_14_ and (CAG)_15_ DNA sequences have observed similar effects on the population of slipped hairpin. ^19^

### Free energy spectrum of the (CAG)*n* polymers as a function of *C*_*S*_

Our theory is that the free energy gap, ∆*G*_*S*_, separating the GS and the excited state, which contains one or more overhangs of CAG repeats at the terminal, is the key determinant of the association rate between RNA chains. To test the theory, we first computed the free energy spectrum for (CAG)_30_ and (CAG)_31_ for 0.15 ≤ *C*_*S*_ ≤ 0.5 M using Eq. 3 (Figure 2 and S2 in SI). As is evident from Fig. 2A and Fig. S2A, ∆*G*_*S*_ separating the GS and the first excited state, with two CAG overhangs, increases with increasing *C*_*S*_. In contrast, the GS of CAG _31_ has one CAG overhang (Fig. 2B), which is susceptible to self-association. In principle, ∆*G*_*S*_ is 0 for (CAG)_31_ as the GS itself contains an overhang region. However, the free energy of the GS increases as *C*_*S*_ increases, which is reflected in the decrease in *P* (*Q*_*HP*_ = 1) (Fig. 1D). As a result, one should expect the dimerization rate for (CAG)_31_ to decrease with an increase in *C*_*S*_. We show ∆*G*_*S*_ and *G*_*S*_ as a function of *C*_*S*_ for (CAG)_30_ and (CAG)_31_, respectively, in Fig. 3A and Fig. 3B. We find that ∆*G*_*S*_ as a function of *C*_*S*_ increases, which suggests that the association of repeat RNAs through homotypic interactions should decrease with an increase in *C*_*S*_.

**Figure 2:**
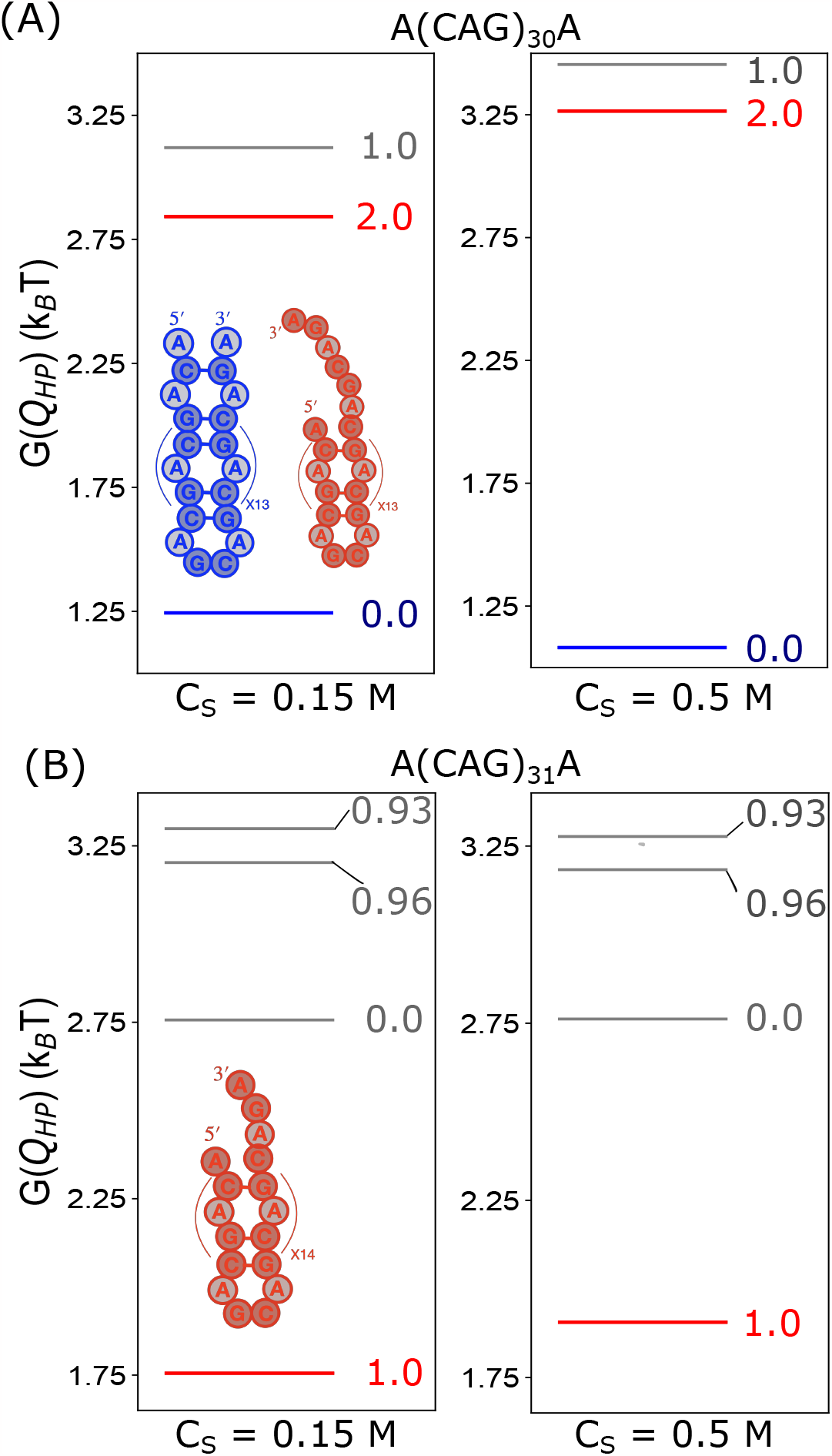
Free energy spectra: (A) *G*(*Q*_*HP*_) as a function of *Q*_*HP*_ at *C*_*s*_ = 0.15 M (left) and 0.5 M (right) for (CAG)_30_. The free energy gap separating the GS and the first excited state with an overhang region increases as *C*_*S*_ increases. A representative GS structure and the first excited state structure are in blue and red, respectively (left panel inset). (B) *G*(*Q*_*HP*_) for (CAG)_31_ at *C*_*s*_ = 0.15 M (left) and 0.5 M (right). The free energy of the GS increase as the *C*_*S*_ increases. The GS conformation (in red in the left panel) has an overhang at the terminal region.

**Figure 3:**
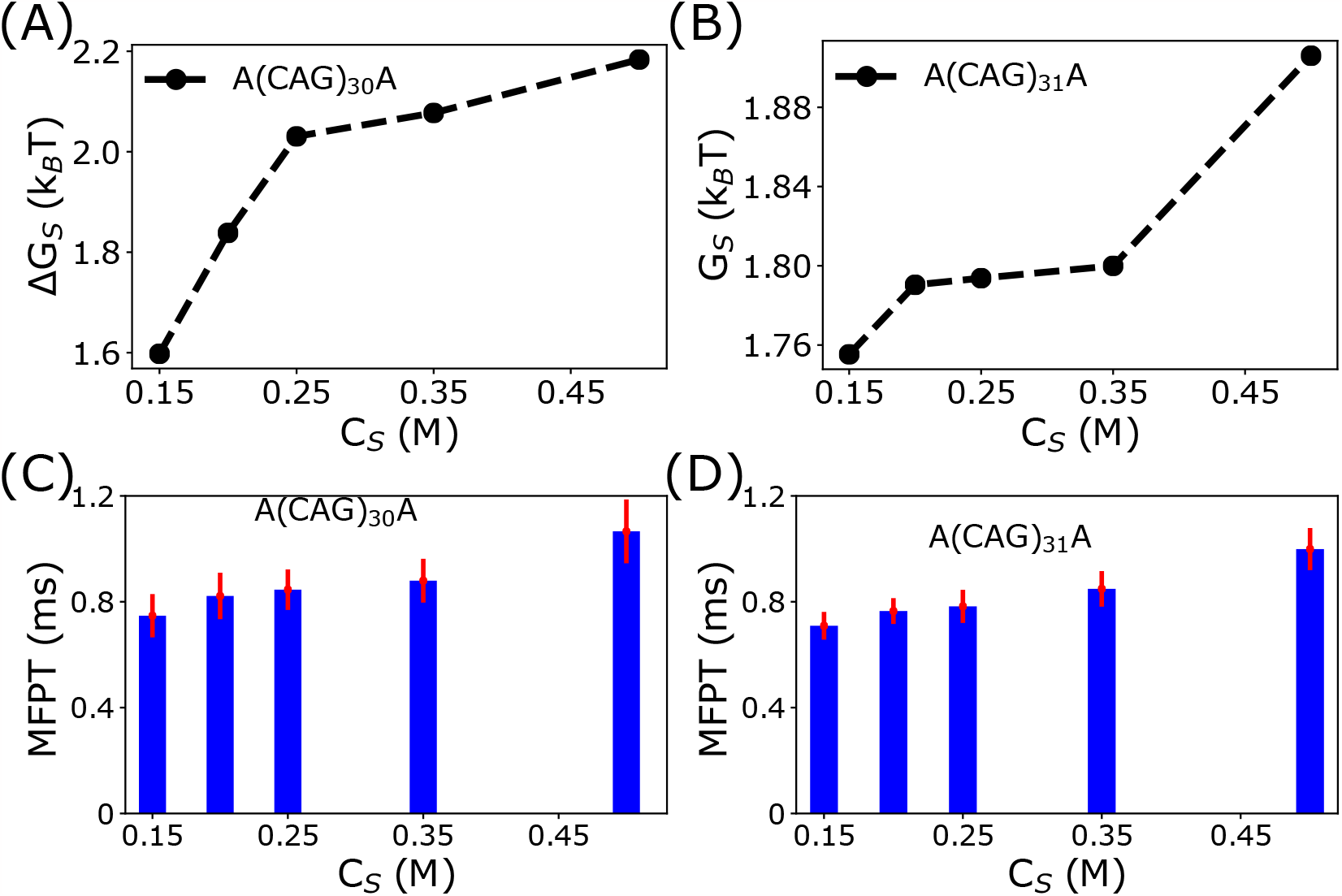
Link between ∆*G*_*S*_ and the mean first passage time, MFPT, for dimerization: (A) ∆*G*_*S*_ as a function of *C*_*s*_ for (CAG)_30_. Increase in ∆*G*_*S*_ as a function of *C*_*S*_ implies that the relative population of aggregation-prone conformation decreases. (B) *G*_*S*_ as a function of *C*_*S*_ is for (CAG)_31_. Increase in *G*_*S*_ with an increase in *C*_*S*_ indicates suppression in population of ground state configurations which are prone to aggregate. (C) The MFPT for (CAG)_30_ at different *C*_*S*_. The standard error in MFPT computed using the bootstrap sampling technique (see Figure S6 in SI) is in red. (D) MFPT for dimerization of (CAG)_31_ increases with an increase in *C*_*S*_, thus correlating with ∆*G*_*S*_.

### Dimerization of hairpins as a function of *C*_*S*_

To test our prediction, we performed Brownian dynamics simulations for dimer formation starting from the ground state of the hairpin structures. The simulations were performed by confining four RNA chains in the hairpin conformation inside a sphere of radius, *R*_0_ (see SI for details). We set *R*_0_ = 100 °A so that initially the monomers interact only weakly with each other. We ran 100 independent trajectories monitoring the formation of dimer for 0.15 ≤ *C*_*S*_ ≤ 0.5 M at 27^*◦*^ C. We calculated the fraction of bps, 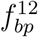, formed between two chains as a function of time to assess if the hairpins formed a duplex (Figure S4). If 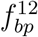 exceeds 0.5, the monomers are in the duplex structure. We computed the number of trajectories leading to the dimer state, *P*_*D*_, as a function of time, *t* (Figure S5). There are ≈ 55 (≈ 42) trajectories in which a dimer formed for (CAG)_31_ ((CAG)_30_) at *C*_*S*_ = 0.15 M. The number of dimer forming trajectories for (CAG)_31_ ((CAG)_30_) decreases to 47 (29) as the *C*_*S*_ is increased to 0.5 M from 0.15 M. In the the concentration range (0.15 *< C*_*S*_ *<* 0.5 M), the number of dimer forming trajectories do not vary significantly.

To compare the propensity to form a dimer at different *C*_*S*_, we calculated the mean first passage time (MFPT), at each salt concentration (Figure 3). The first passage time (FPT), *τ*_*F P T*_, for each dimer forming trajectory is identified with the time at which 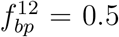. The MFPT is given by 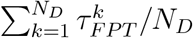, where, 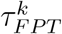 is the FPT for the *k*^*th*^ trajectory and *N*_*D*_ is the total number of dimer forming trajectories. Fig. 3C and 3D show the MFPT as a function of *C*_*S*_. Our finding that the rate of dimerization (∝ MFPT^*−*1^) decreases as *C*_*S*_ increases accords well with the theoretical prediction. The modest change in MFPT in varying *C*_*S*_ is related to the small change in ∆*G*_*S*_ as a function of *C*_*S*_ in the wild type sequences.

### Hairpin opening is the rate determining step in RNA aggregation

Salts influence the time (*τ*_*conv*_) required for unwinding the intra-molecular bps during the conversion of hairpins into an anti-parallel duplex structure by modulating the stability of intra-molecular bps. To investigate the effects of salts on *τ*_*conv*_, we computed *τ*_*conv*_ for each trajectory at *C*_*S*_ = 0.15 M and 0.5 M. Consider the part of the trajectory which leads to the formation of dimer (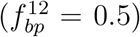) without re-entering into the hairpin state. We define *τ*_*conv*_ as the time required for dimerization to be complete once association between the chains starts (Figure S7A). *τ*_*conv*_ for the dimer forming trajectories at *C*_*S*_ = 0.5 M is considerably longer compared to *C*_*S*_ = 0.15 M, which is also reflected in the average ⟨*τ*_*conv*_⟩ values. The ratio of *τ*_*conv*_ obtained at *C*_*S*_ = 0.5 M to 0.15 M is ≈ 2.2 for (CAG)_30_ and ≈ 2.3 for (CAG)_31_ (Figure S7B and S7C). Our results are in accordance with the experimental observation^21,22^ that the hairpin opening rate strongly depends on the ionic concentrations.

### Reduction in strand slippage in scrambled sequences decreases the self-association propensity

In order to further illustrate the link between phase behavior of RNA to the monomer characteristics, we calculated the free energy spectra for scrambled sequences whose sequence lengths and composition are the same as (CAG)_31_ but the nucleotide positions are shuffled. We chose two sequences SS1 and SS2. The sequences are given in Figure 4. The free energy spectra at *C*_*S*_ = 0.15 M and 0.5 M are given in Figure 4 and Figure S8, respectively. In contrast to the GS of (CAG)_31_, which has one CAG overhang at the terminal, the GS of the scrambled sequences do not have overhangs. The value of *Q*_*HP*_ is zero for the GS of both SS1 and SS2. The GS conformations for the sequences are shown in Figure 4B and Figure S9. The terminal nucleotides of the scrambled sequences are engaged in the formation of intramolecular base pairing in the GS conformations, and therefore cannot easily self-associate. However, the RNA chains may aggregate by accessing the excited state with overhangs. The excited states of the SS1 are devoid of such structures because of the consecutive array of GC base pair formation at the terminal. We surmise that the propensities of SS1 and SS2 to self-associate are small.

**Figure 4:**
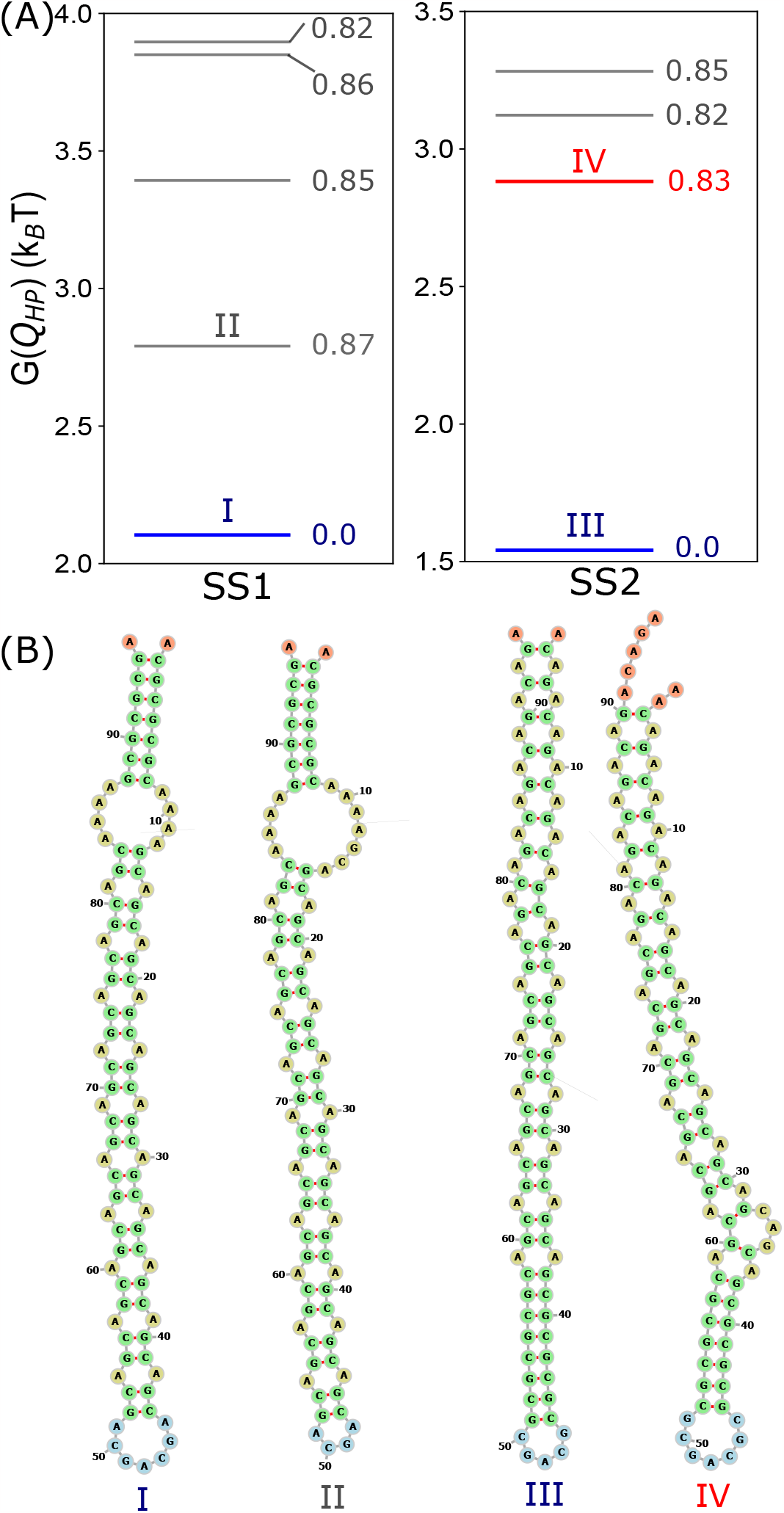
Free energy spectra for scrambled sequences at *C*_*S*_ = 0.15 M: (A)*G*(*Q*_*HP*_), as a function of *Q*_*HP*_ for the sequence SS1 (left panel). The SS1 sequence is A(CG)_3_CA_4_G(CAG)_23_CA_4_G(CG)_3_A. The GS and the excited states are in blue and gray lines, respectively. There is no overhang at the terminal, which is required for self-association. Right panel shows *G*(*Q*_*HP*_), as a function of *Q*_*HP*_, for SS2 whose sequence is (ACAG)_4_(CAG)_7_(CG)_4_CAG(CG)_4_(CAG)_7_(ACAG)_4_A. The GS with no overhang and the first excited state with an overhang at the end are in blue and red lines, respectively. (B) GS and the first excited state conformations of the hairpin structures for SS1 (I and II) and SS2 (III and IV).

The absence of multiple peaks in the end-to-end distance distribution (*P* (*R*_*ee*_)) further supports our conclusion that the sequence SS1 contains overhang(s) neither in the GS nor in the excited states (Figure S10A). Based on the free energy spectrum, we predict that the sequence SS1 has the lowest tendency to undergo self-association. In contrast, the first excited state of the SS2 contains an overhang at the terminal of the hairpin structure (Figure 4B and S10B). The SS2 sequence is, therefore, susceptible to the formation of higher-order oligomeric structures. However, the propensity to form the oligomers is less compared to the (CAG)_31_ because of enhancement in the stability of the GS for the scrambled sequence (Figure S8). The enhanced stability is also reflected in the distribution of the end-to-end distance (Figure S10B). The *P* (*R*_*ee*_) value at *R*_*ee*_ ≈ 1.45 nm increases significantly as *C*_*S*_ increases from 0.5 M from 0.15 M. The value of ∆*G*_*S*_ also increases as *C*_*S*_ increases. Because the rate of dimer formation correlates inversely with the free energy gap, we predict that the formation of higher-order oligomeric structure in SS2 should decrease substantially as the *C*_*S*_ is increased relative to the wild-type low complexity sequence.

Using the scrambled sequence as a reference, both experiments^1^ and simulations^14^ investigated the effects of nucleotide position in the sequences in the context of phase separations of CAG repeat RNAs. It was shown there is a suppression in the propensity for aggregation in the scrambled sequences, which is due to the increase in ∆*G*_*S*_. Shuffling of the nucleotide positions in the sequence also changes the nature of the free energy spectrum, thus modulating ∆*G*_*S*_, which is the single most important determinant of the phase behavior of RNA sequences.

## Conclusions

We established a direct link between changes in the monovalent salt concentration on the free energy spectrum of low complexity RNA sequences and their propensities for self-association. By combining counterion condensation theory, with Debye-Huckel potential for the electrostatic interactions, we accounted for the effects of monovalent salts in coarse-grained SIS model^23^ for RNA. The major finding of our study is that the population of the hairpin state, corresponding to self-association prone conformations, is suppressed for both (CAG)_30_ and (CAG)_31_ as *C*_*S*_ is increased. The free energy gap, ∆*G*_*S*_, separating an aggregation-prone hairpin state from an aggregation inactive state, increases with an increase in *C*_*S*_. Strikingly, the mean first passage time (MFPT) for dimer formation increases with ∆*G*_*S*_. More generally, our theory suggests that there is anti-correlation between ∆*G*_*S*_ and rate of RNA association. Because ∆*G*_*S*_ for a given sequence can be altered by changing external conditions, we predict that condensate formation may be drastically changed by tuning temperature and crowding. Our result that the dimer formation rate decreases as the salt concentration is increased can be tested experimentally.

Interestingly, the calculated values of ∆*G*_*S*_ for RNA repeat sequences are similar to the experimentally inferred values for the corresponding DNA sequences. For example, from the population of low and high FRET states of CAG_14_ and CAG_15_ at low salt concentration (see Fig. 2 in Ref.^19^), we find that ∆*G*_*S*_ ≈ 1.45 *k*_*B*_*T* . In the RNA repeat sequences value of ∆*G*_*S*_ is ≈ (1.3 − 2.3) *k*_*B*_*T* at a higher value of the salt concentration. Just like the DNA trinucleotide repeats the corresponding RNA sequences are dynamic, which results in multiple states being sampled. It would be valuable to perform single molecule FRET experiments for low complexity RNA sequences in order to verify some of our findings.

An important finding in our study is that ∆*G*_*S*_ increases substantially for the scrambled sequences relative to the repeat sequences. Therefore, we expect that the time for dimerization for the scrambled sequences should be substantially greater for the scrambled sequences. Assuming that the MFPT ∝ *exp*(−∆*G*_*S*_*/k*_*B*_*T*),^24^ we predict that the time for SS2 to dimerize should be roughly ten times greater. Because ∆*G*_*S*_ depends on the precise sequence it follows that from the astronomically large number of sequences (= 3^*n*^) it is possible to construct sequences that would not form condensates at any reasonable external conditions. Thus, salt-dependent ∆*G*_*S*_ may be used to control aggregation of RNA.

Our predictions are significant in the cellular context. Physiologically relevant ionic concentration varies between cells of different species. For instance, the ionic concentration of potassium ion (K^+^) in budding yeast can reach up to 300 mM^25^ whereas it is ≈ 150 mM in mammalian cells. The free energy spectrum generated for a single RNA molecule in the presence of different ionic concentrations could be used as an important tool to regulate the phase behavior of RNA-rich condensates.

## Methods

### Models

Following our earlier studies, ^14,15,23^ we represent each nucleotide by a single bead. In order to account for counter ion condensation effects, ^26,27^ we used a reduced value of charge -*Q* (0 *< Q <* 1) on the bead. In monovalent salt solutions 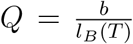 where, *b* = 4.4 °A,^28^ and 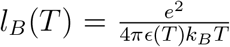, is the Bjerrum length with *e* being the electron charge, *k*_*B*_ is the Boltzmann constant, and T is the temperature. The *T* dependence of the dielectric constant^29^ is *ϵ*(*T*) = 87.74 - 0.4008*T* + 9.398 × 10^*−*4^ *T* ^2^ - 1.410 × 10^*−*6^ *T* ^3^, and, T is expressed in ^*◦*^C unit. The electrostatic repulsion between the beads, accounting for the interactions between the phosphate groups, is given by,

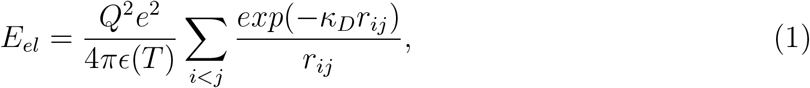

where *κ*_*D*_ = (8*πρl*_*B*_)^1*/*2^, *ρ* is the number density of the monovalent ions. The total energy in the Single Interaction Site (SIS) model is, *E*_*T OT*_ = *E*_*B*_ + *E*_*HB*_ + *E*_*EV*_ + *E*_*el*_. The detailed functional forms of the bonded (*E*_*B*_), hydrogen bond (*E*_*HB*_), and the excluded volume interactions (*E*_*EV*_) along with the parameter values are given elsewhere. ^15^ Although the SIS model is simple, our previous study^15^ showed that the agreement between the calculated and experimentally measured heat capacities for several (CAG)_n_ constructs is very good.

### Free energy spectra

We first calculated the distribution, *P* (*Q*_*HP*_), of the hairpin order parameter, *Q*_*HP*_, which measures the deviation of WC base pairs (GC base pairs in our case) with respect to a perfectly aligned hairpin structure. The *Q*_*HP*_ order parameter is defined as,

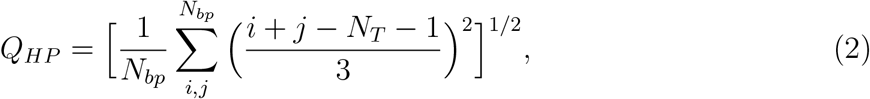

where *i* and *j* are the nucleotides, *N*_*T*_ is the sequence length, and *N*_*bp*_ is the number of base pairs in a given conformation. For a perfect hairpin, it is easy to show that *i* + *j* = *N*_*T*_ + 1, implying that *Q*_*HP*_ = 0. The value of *Q*_*HP*_ = 1 corresponds to a slipped hairpin, corresponding to one unit of unpaired CAG. Fractional values, 0 *< Q*_*HP*_ *<* 1, represent a variety of conformations containing bulges in the stem. The free energies are calculated by arranging *P* (*Q*_*HP*_) values in descending order. The spectrum is computed using,

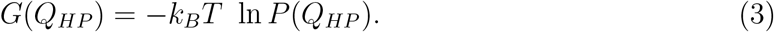

### Simulations

The thermodynamic properties, including the free energy spectra, are calculated using the trajectories generated by integrating the Langevin equation of motion in the low friction limit. ^30^ In order to investigate the formation of aggregates in (CAG)_30_ and (CAG)_31_, we performed Brownian dynamics simulations using the Ermack-McCammon algorithm.^31^ The details are given in the Supplementary Information.

## Acknowledgements

This work was supported by a grant from the National Science Foundation (CHE 2320256) and the Welch Foundation through the Collie-Welch Chair (F-0019).

## TOC figure

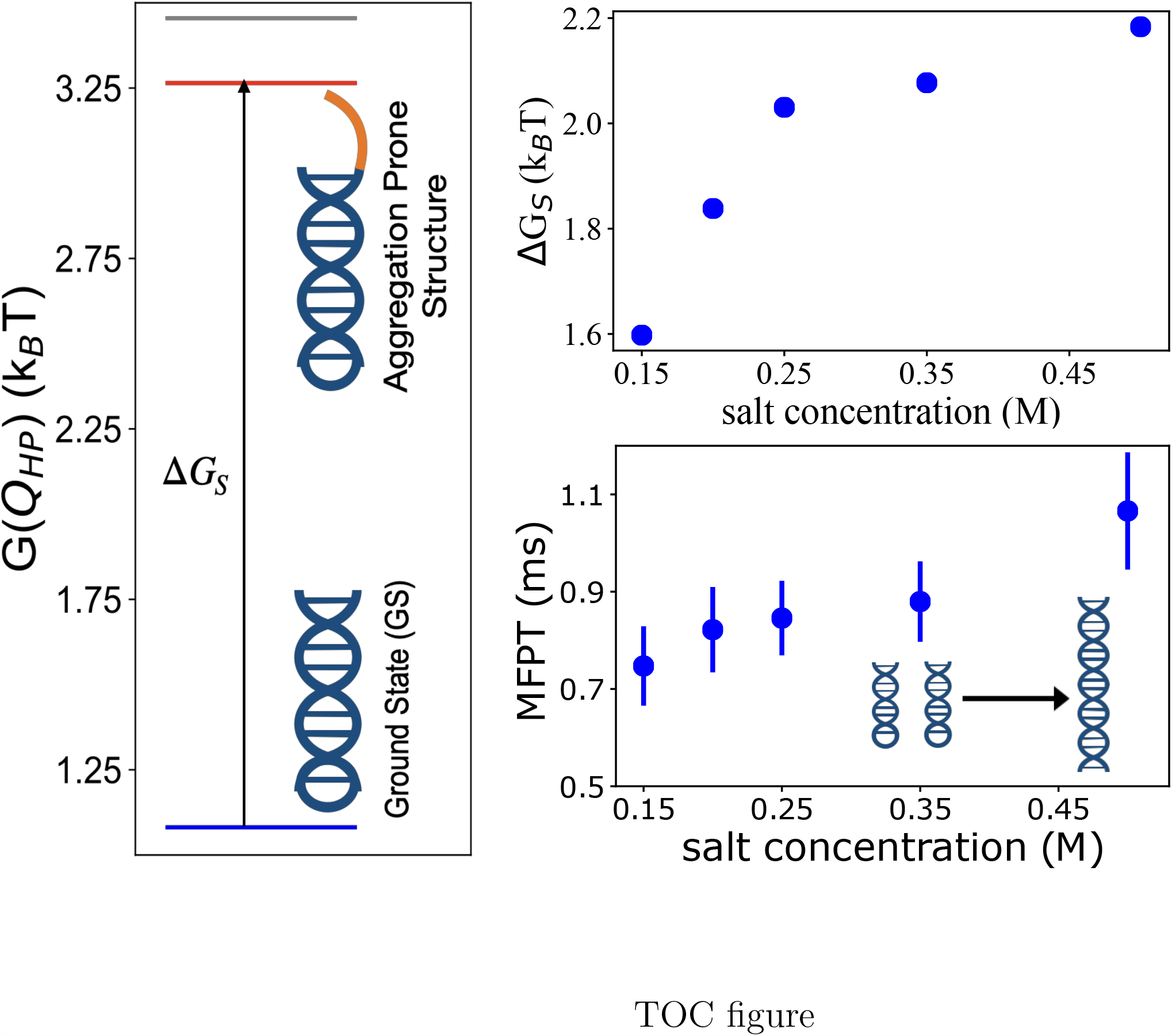

## Notes

### Competing Interest Statement

The authors have declared no competing interest.

