## Supporting Information for "Salt-dependent self-association of trinucleotide repeat RNA sequences"

‖

**Model:** In the coarse-grained (CG) Self-Organized Polymer (SOP) model RNA, each nucleotide is in the RNA sequence is represented by a single bead, as in the Self-Organized Polymer (SOP) model.[1] The total energy, $E_{TOT}$, is the sum of bonded ($E_B$), hydrogen bonded ($E_{HB}$), excluded volume ($E_{EV}$), and electrostatic energy ($E_{el}$), and is written as,

$$E_{TOT} = E_B + E_{HB} + E_{EV} + E_{el}. \tag{S1}$$

In Eq. S1, $E_B$ is the sum of bond stretch ($E_{BS}$) and bond angle ($E_{BA}$) potentials. The expression for $E_B$ is,

$$E_B = E_{BS} + E_{BA} \equiv \frac{1}{2}\sum_{i}^{N_T-1} k_{bond}(r_i - r_0)^2 + \frac{1}{2}\sum_{j}^{N_T-2} k_{angle}(\alpha_j - \alpha_0)^2, \tag{S2}$$

where $N_T$ is the number of nucleotides in a given sequence. The parameters, $k_{bond}$ and $k_{angle}$, are the spring constants in $E_{BS}$ and $E_{BA}$, respectively. In Eq. S2, $r_i$ and $\alpha_j$ are the $i^{th}$ bond distance and $j^{th}$ bond angle, respectively, and $r_0$ and $\alpha_0$ are the corresponding equilibrium values.

Hydrogen bond interactions between two beads, with complementary base pairs such as cytosine (C) and guanine (G), and, adenine (A) and uracil (U), contribute substantially to the stability of the folded hairpin-like structures of the low complexity sequences. The functional form for $E_{HB}$ is,

$$E_{HB} = N_b u_{hb}^0 \exp[-U_{hb,bond} - U_{hb,angle} - U_{hb,dihedral}], \tag{S3}$$

where $N_b = 3$ (2) for a G-C (A-U) base pair (bp), and $u_{hb}^0$ (= -2 kcal/mol) is the strength of a single hydrogen bond.

The expressions for $U_{hb,bond}$, $U_{hb,angle}$ and $U_{hb,dihedral}$ are,

$$U_{hb,bond} = k_r(r_{ij} - r_{hb,0})^2, \tag{S4}$$

$$U_{hb,angle} = k_\theta(\theta_{i,j,j-1} - \theta_1)^2 + k_\theta(\theta_{i-1,i,j} - \theta_1)^2 + k_\theta(\theta_{i,j,j+1} - \theta_2)^2 + k_\theta(\theta_{i+1,i,j} - \theta_2)^2, \quad \text{(S5)}$$

$$U_{hb,dihedral} = k_\phi[1 + cos(\phi_{j-1,j,i,i-1} + \phi_1)] + k_\phi[1 + cos(\phi_{j+1,j,i,i+1} + \phi_2)]. \quad \text{(S6)}$$

In Eq. S4, $r_{ij}$ is the distance between beads $i$ and $j$, $r_{hb,0}$ is the hydrogen bond distance in an ideal A-form helix, and $k_r$ is a constant. In Eq. S5, $\theta_{i,j,k}$ is the angle between beads $i$, $j$ and $k$, and $\theta_1$ and $\theta_2$ are two constants. Contribution of the angular part to the total hydrogen bond potential is determined by $k_\theta$, which has the unit of radian$^{-2}$. In Eq. S6, the dihedral angle, $\phi_{i,j,k,l}$, is the angle between the two planes defined by the four beads $i$, $j$, $k$ and $l$, and, $k_\phi$, $\phi_1$ and $\phi_2$ are constants.

In order to avoid unphysical overlap between two non-bonded beads that are separated by two other beads, we used a repulsive potential,

$$E_{EV} = \epsilon_{ev}[(\frac{\sigma}{r_{ij}})^{12} - (\frac{\sigma}{r_{ij}})^6 + 1] \text{ if } r_{ij} \leq \sigma, \quad \text{(S7)}$$

$$E_{EV} = 0 \text{ otherwise}, \quad \text{(S8)}$$

where, $\epsilon_{ev}$ is the strength of repulsive interaction, and $\sigma = 10$ Å. The values of the parameters in the energy function are listed in Table S1.

**Electrostatic interactions and charge renormalization:** In the presence of cations, the charge on each bead is reduced to -Q ($0 < Q < 1$) from -1 due to counterion condensation.[2,3] The effects of monovalent salt ions are included implicitly in the SOP model by using a renormalized $Q$ value. We calculated $Q$, in presence of monovalent salts, as $\frac{b}{l_B(T)}$, where $b$ (=4.4 Å)[4] is the length per unit charge. The Bjerrum length, $l_B(T) = \frac{e^2}{4\pi\epsilon(T)k_BT}$ where $e$ is the electron charge, $\epsilon(T)$ is the temperature dependent dielectric constant of water, $k_B$ is the Boltzmann constant, and $T$ is the temperature. The temperature dependent $\epsilon(T)$, $T$ (in °C) is computed using,[5]

$$\epsilon(T) = 87.74 - 0.4008T + 9.398 * 10^{-4}T^2 - 1.410 * 10^{-6}T^3. \quad \text{(S9)}$$

The electrostatic energy between two negatively charged phosphate groups is taken to be,

$$E_{el} = \frac{Q^2 e^2}{4\pi\epsilon(T)} \sum_{i<j} \frac{\exp(-\kappa_D r_{ij})}{r_{ij}}, \tag{S10}$$

where $\kappa_D$ ($=\sqrt{8\pi\rho l_B(T)}$) is the inverse Debye length. In monovalent salt solutions $q_i = 1$ is the magnitude of the charge on each counterion, and $\rho$ is the number density of ions in the solution. For a given monovalent salt concentration, $C_s$, it is easy to show that $\rho = 2C_s \times 6.022 \times 10^{-4}$ Å$^{-3}$.

**Inter-chain interactions in dimers:** Interactions between nucleotides belonging to two RNA chains are the same as in intra-chain interactions. In order to maintain proximity between the four chains to ensure that molecular association occurs on a reasonable time scale in the simulations, the center of mass of the chains are constrained by a harmonic potential, $U_D$,[6]

$$U_D = \frac{1}{2} k_D (R_{cm} - R_0)^2 \text{ if } R_{cm} > R_0. \tag{S11}$$

If $R_{cm} < R_0$ then we set $U_D = 0$ . In Eq.S11, $R_{cm}$ is the distance between the center of mass of a chain and the center of the sphere, and, $R_0$ is the radii of the sphere. We used $R_0 =$ 100 Å . The value of $k_D$ is 200 kcal/(mol.Å$^2$). Note that for the restraining potential in Eq. S11 the precise value of $R_0$ has negligible effect on dimer formation rates or yields.

**Simulations:** To compute the equilibrium thermodynamic properties of $(\mathrm{CAG})_\mathrm{n}$ monomers, we performed low friction Langevin dynamics simulations[7] by integrating the equation of motion, which for bead $i$ is,

$$m\ddot{\vec{r_i}} = -\zeta\dot{\vec{r_i}} + \vec{F_C} + \vec{\Gamma}. \tag{S12}$$

In the above equation, $m$ is the mass of each bead, $\zeta$ is the friction coefficient, $F_C = -\frac{\partial E_{TOT}}{\partial \vec{r_i}}$, and $\vec{\Gamma}$ is the random force with Gaussian statistics, $\langle \Gamma(t)\, \Gamma(t+ph) \rangle = \frac{2\zeta k_B T}{h} \delta_{0,p}$ where $p = 0, 1, \cdots$, $\delta_{0,p}$ is the Kronecker delta function. We integrated Eq. S12 using the velocity Verlet algorithm. Following our previous work,[7] we used a small value of the friction coefficient in order to sample the conformational space efficiently. The friction coefficient $\zeta = 0.05\ \frac{m}{\tau_L}$ and $h = 0.005\ \tau_L$, where $\tau_L$ $(= \sqrt{\frac{m_0 a_0^2}{\epsilon_h}})$ is the time unit in the Langevin dynamics simulations. Thermodynamic properties do not depend on the precise value of $\zeta$ or $m$.

To investigate the kinetics of dimer formation, we performed Brownian dynamics simulations using the Ermak-McCammon algorithm,[8]

$$\dot{\vec{r_i}}(t+h) = \vec{r_i}(t) + \frac{h}{\zeta}\vec{F_C}(t) + \vec{R}(t), \tag{S13}$$

where $\vec{R}(t)$ is the random displacement with mean $\langle R(h)\rangle = 0$, and variance $\langle [R(h)]^2\rangle = \frac{2k_BTh}{\zeta}$. The time step, $h$, to integrate the equation of motion is taken as $0.1\tau_H$, where, $\tau_H$ is the time unit of Brownian dynamics simulations and $\zeta$ is taken as 26.6 $m/\tau_H$. The value of $\tau_H = \frac{\zeta_H a_0^2}{k_BT} \approx 400$ ps.

**Generation of scrambled sequences:** Experiments[9,10] and simulations[11] showed that precise ordering of the multivalent interacting sites in the sequence is crucial in determining the aggregation propensity. To demonstrate the role of sequence effects simulations[11] and experiments[9] used scrambled sequences as a reference system. In scrambled sequences, the chain length and the sequence composition remain the same as in the wild-type sequence, but the position of the nucleotides is shuffled arbitrarily. For a trinucleotide repeat sequence such as $(CAG)_n$, $3^n$ number of sequences are realizable in principle but the number would be smaller if the sequence composition is fixed. To demonstrate how the free energy spectrum of the scrambled sequences varies with respect to wild-type sequences, we generated two sequences by fixing the composition and the sequence length to be the same as $(CAG)_{31}$. During the selection procedure of the scrambled sequences, we emphasized whether the scrambled sequences enrich or deplete the CG base pairs at the terminal regions. We found SS1 (sequence: $A(CG)_3CA_4G(CAG)_{23}CA_4G(CG)_3A$) enriched the CG base pairs at the terminal whereas SS2 (sequence: $(ACAG)_4(CAG)_7(CG)_4CAG(CG)_4(CAG)_7(ACAG)_4A$)

deplete the CG base pairs at the terminal. There are large number of ways of generating scrambled sequences. All of them would enhance sequence heterogeneity, and hence would suppress their propensity to self-associate.

**Table S1: List of parameters used in the single bead SOP energy function**

| | |
|---|---|
| $k_{bond}$ | 15 kcal/mol. Å$^{-2}$ |
| $r_0$ | 5.9 Å |
| $k_{angle}$ | 10 kcal/mol.rad$^{-2}$ |
| $\alpha_0$ | 2.618 rad |
| $\epsilon_{EV}$ | 2.0 kcal/mol |
| $\sigma$ | 10 Å |
| $r_{hb,0}$ | 13.8 Å |
| $k_r$ | 3.0 Å$^{-2}$ |
| $k_\theta$ | 1.5 rad $^{-2}$ |
| $k_\phi$ | 0.5 |
| $\theta_1$ | 1.8326 rad |
| $\theta_2$ | 0.9425 rad |
| $\phi_1$ | 1.8326 rad |
| $\phi_2$ | 1.345 rad |
| $u_{hb,0}$ | -2.0 kcal/mol |

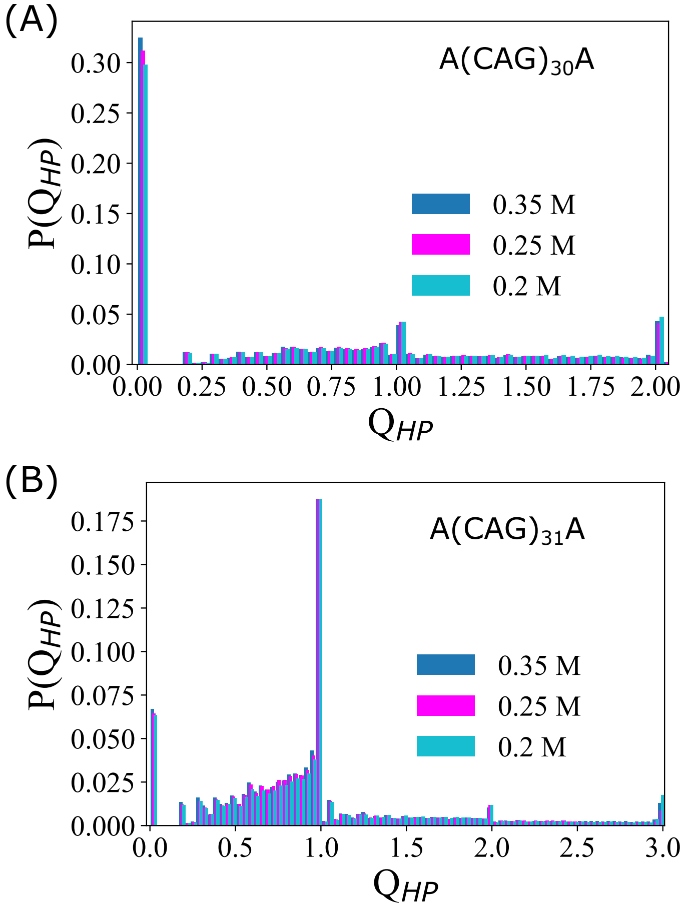


Figure S1: Probability $P(Q_{HP})$ distribution of the order parameter $Q_{HP}$ order parameter: (A) $P(Q_{HP})$ for $(\text{CAG})_{30}$ at salt concentration, $C_s = 0.35$ M, 0.25 and 0.2 M are in blue, magenta and cyan, respectively. With an increase in $C_s$ the probability of $Q_{HP} = 0$ increases whereas $P(Q_{HP})$ with $Q_{HP} > 0$ decreases. (B) $P(Q_{HP})$ for $(\text{CAG})_{31}$ at $C_s = 0.35$ M (blue), 0.25 (magenta) and 0.2 M (cyan) indicates the ground state population of $(\text{CAG})_{31}$ decreases, i.e. $P(Q_{HP})$ at $Q_{HP} = 1$ decreases.

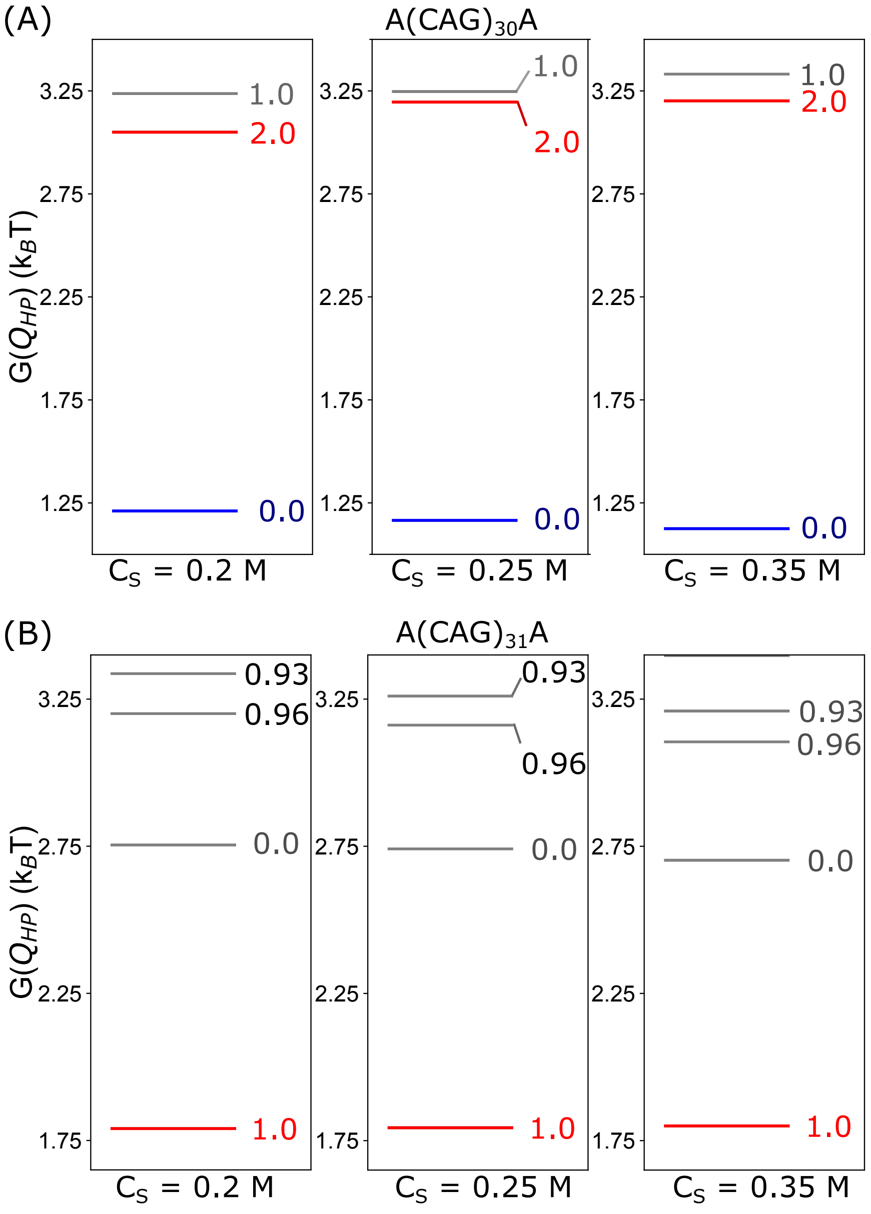


Figure S2: Free energy spectrum: (A) Free energy as a function of $Q_{HP}$ value, $G(Q_{HP})$ for $(CAG)_{30}$, at $C_s$ = 0.20 M (left), 0.25 M (middle) and 0.5 M (right). The free energy gap separating the ground state and the first excited state containing an overhang region increases with an increase in $C_S$. (B) $G(Q_{HP})$ for $(CAG)_{31}$ at $C_s$ = 0.2 M (left), 0.25 M (middle) and 0.5 M (right). The ground state conformation contains an overhang region, making it susceptible to form aggregates with a higher rate.

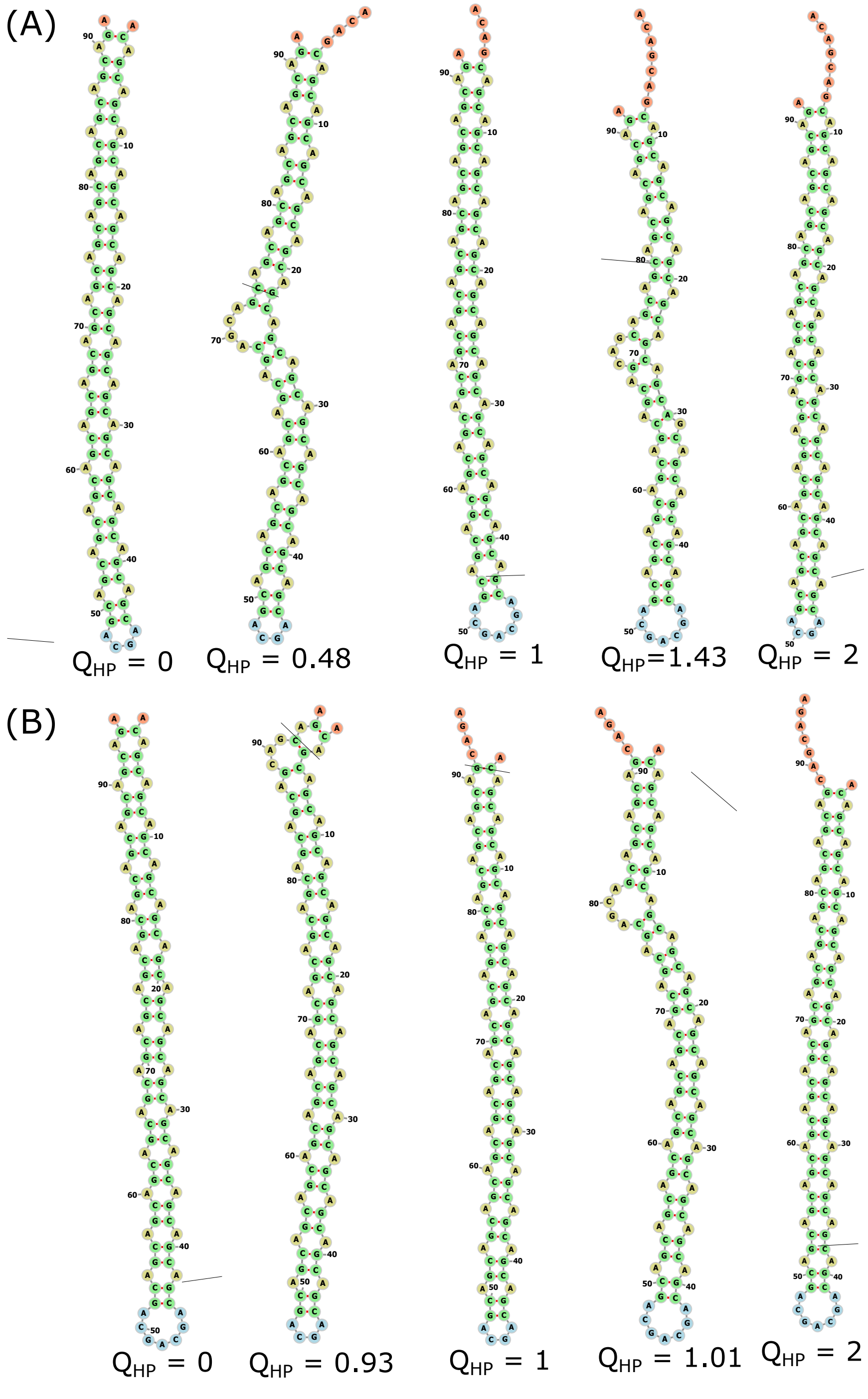


Figure S3: Hairpin structures of $(CAG)_{30}$ and $(CAG)_{31}$: (A) Different conformations of $(CAG)_{30}$ hairpins with their corresponding $Q_{HP}$ values. A hairpin with $Q_H = m$, where, $m = 0, 1, 2...$, contains an overhang region at the terminal region. Hairpins with fractional $Q_{HP}$ values contains a bulge in the stem of the hairpin. The ground state (GS) structure of the $(CAG)_{30}$ has $Q_{HP} = 0$. (B) Hairpin structures of $(CAG)_{31}$ with various $Q_{HP}$ values. The GS structure of $(CAG)_{31}$ has $Q_{HP} = 1$.

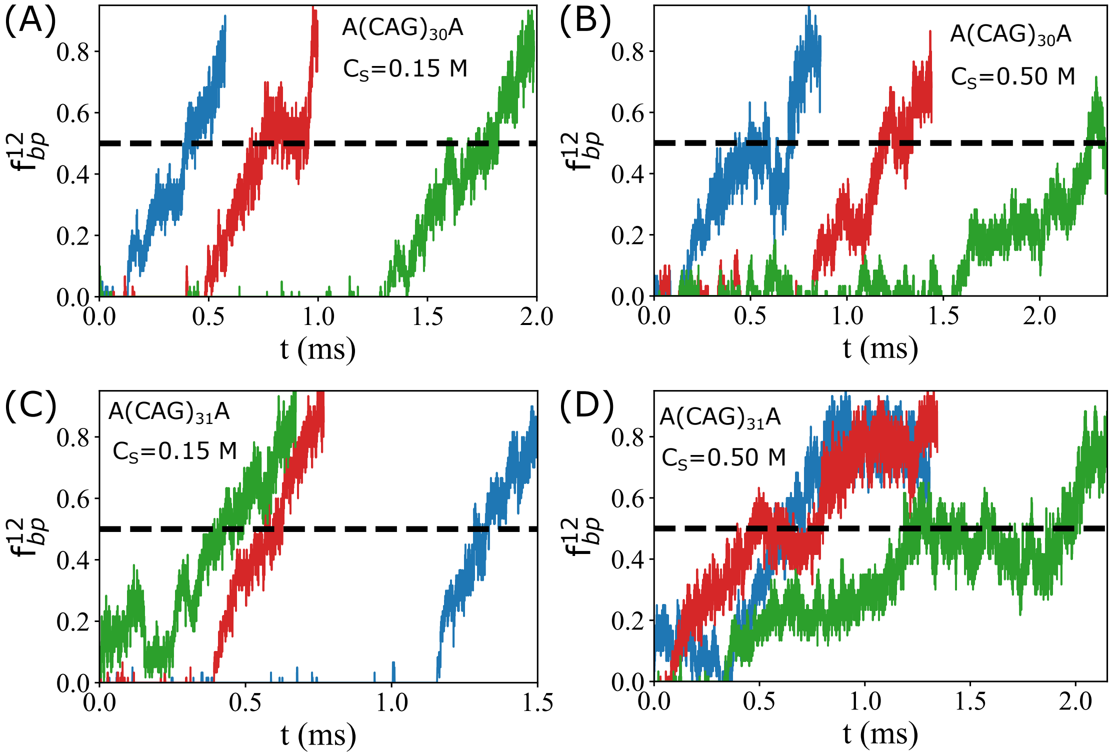


Figure S4: Representative dimer forming trajectories starting from GS structures: Fraction of base pairs, $f_{bp}^{12}$ formed between two monomers as a function of time is computed to monitor the formation of dimer. The black dashed line indicates $f_{bp}^{12} = 0.5$. At each salt concentration, three representative trajectories in red, green and blue leading to the dimer structure are shown. $f_{bp}$ as a function of time for (A) $(CAG)_{30}$ at $C_S = 0.15$ M, (B) $(CAG)_{30}$ at $C_S = 0.5$ M, (C) $(CAG)_{31}$ at $C_S = 0.15$ M, and, (D) $(CAG)_{31}$ at $C_S = 0.5$ M.

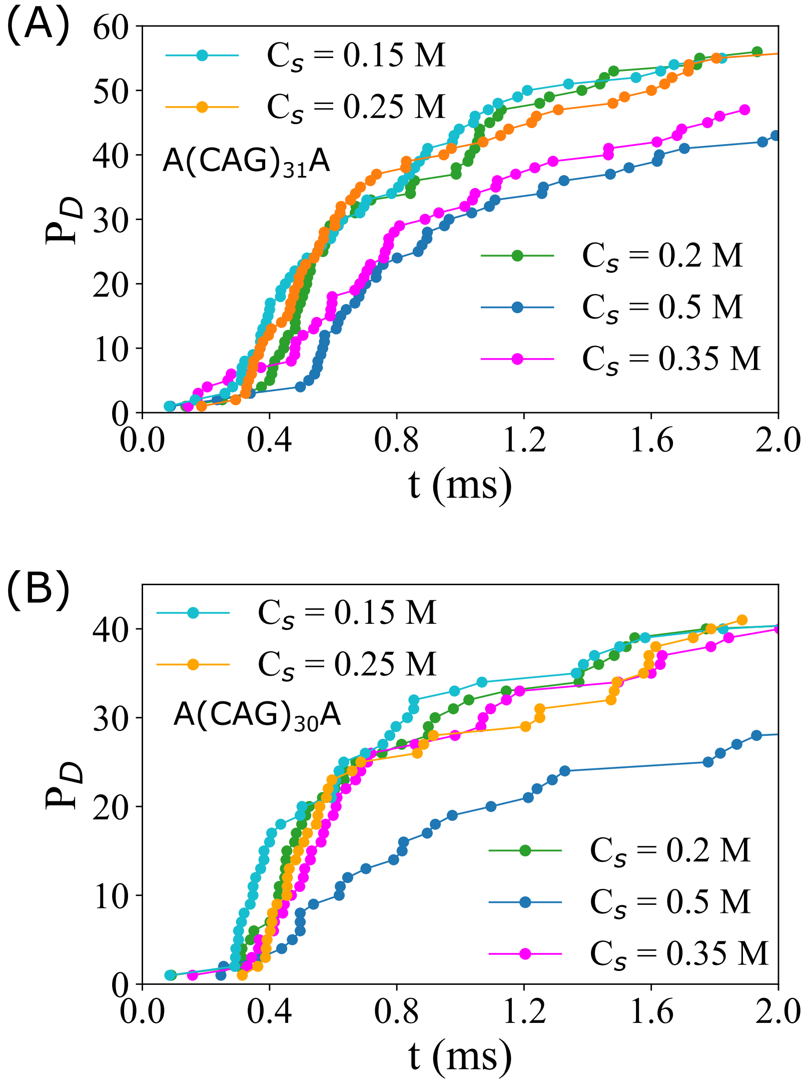


Figure S5: Dimer yield as a function of time: (A) The number of trajectories reaching the dimer state, $P_D$ as a function of time, $t$ for $(CAG)_{31}$ for salt concentration, $C_S = 0.15$ (cyan), 0.2 (green), 0.25 (orange), 0.35 (magenta) and 0.5 M (blue). As $C_S$ is changed from 0.15 to 0.5 M, value of $P_D$ is decreased. (B) Same as (A) except for $(CAG)_{30}$.

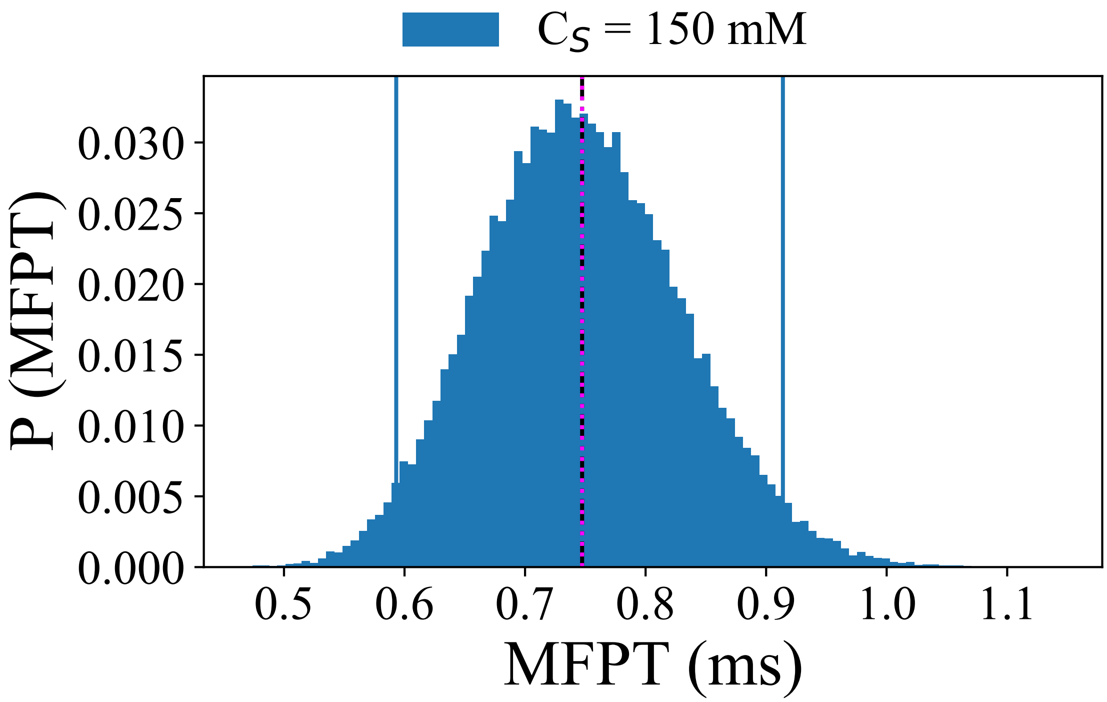


Figure S6: Distribution of the mean first passage time, $P(MFPT)$ for $(\text{CAG})_{31}$, obtained using bootstraping sampling technique. The original MFPT value computed using the actual data set of first passage time is in red dotted line, where as the mean value of MFPT obtained in the bootstarp sampling methods is in black dotted line. The distribution is computed using 50,000 resampled data data set. The 95% confidence interval is shown by the cyan vertical lines. The P(MFPT) is computed using the data set obtained for $C_S = 0.15$ M.

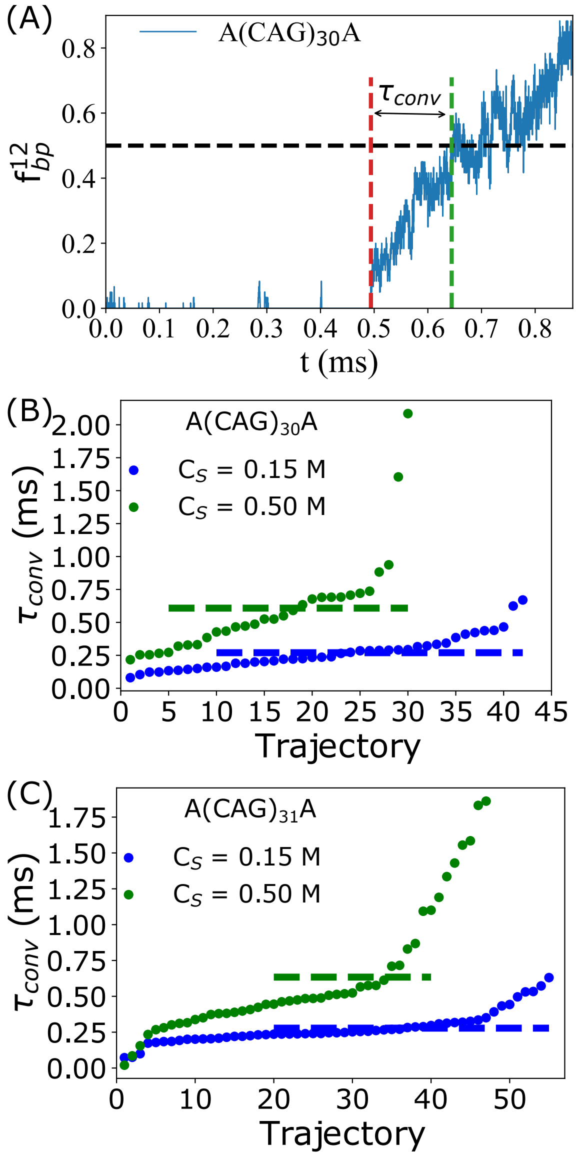


Figure S7: Time required for the completion of a dimer starting from initiation time, $\tau_{conv}$: (A) Fraction of base pairs ($f_{bp}^{12}$) between two monomers of $(CAG)_{30}$ hairpin as a function time. The initiation of the dimer formation is indicated by the red dashed line and completion of the dimer formation is shown by the green line. Time taken to go from initiation to completion point is defined as $\tau_{conv}$, which is indicated by the black arrow. (B) $\tau_{conv}$ obtained for different trajectories leading to dimers from $(CAG)_{30}$ hairpins are in blue and green for $C_S = 0.15$ M and 0.5 M, respectively. The average value of $\tau_{conv}$ ($\langle\tau_{conv}\rangle$) computed at $C_S = 0.15$ M and 0.5 M is in blue dashed line and green dashed line, respectively. (C) Same as (B) except for $(CAG)_{31}$.

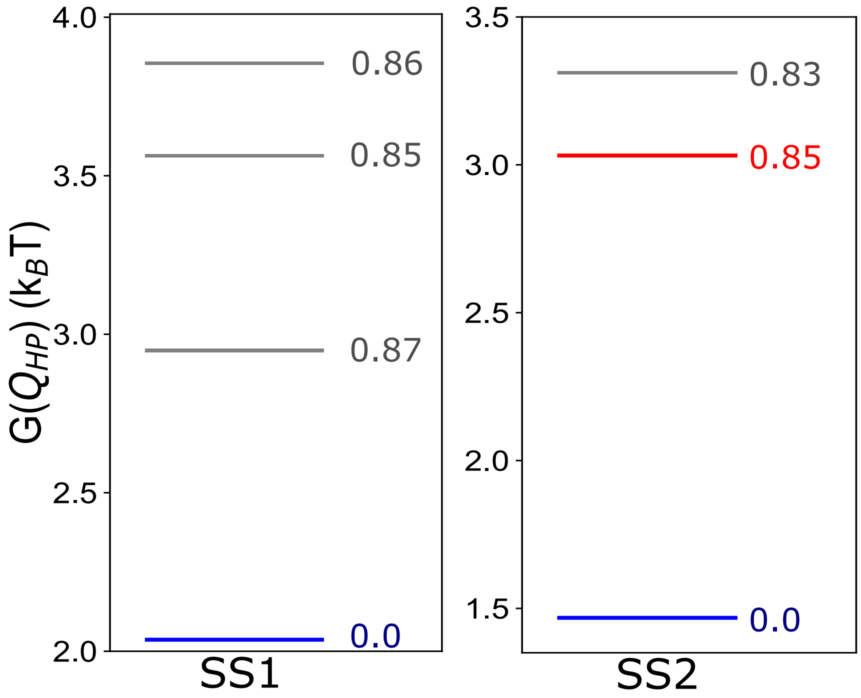


Figure S8: Free energy spectra of scrambled sequences generated at $C_S = 0.5$ M: (right panel) Free energy, $G(Q_{HP})$, as a function of $Q_{HP}$ for the scrambled sequence 1 (SS1). The SS1 sequence is $A(CG)_3CA_4G(CAG)_{23}CA_4G(CG)_3A$. Because of the presence of consecutive CGs base pairs at the terminal region, the GS (in blue) and the excited states do not contain overhangs. Hence, the propensity of SS1 to aggregate should be small. (left panel) $G(Q_{HP})$ for SS2 with sequence, $(ACAG)_4(CAG)_7(CG)_4CAG(CG)_4(CAG)_7(ACAG)_4A$. The GS and the first excited state of the spectrum are in blue and red, respectively. The first excited state of the RNA contains unpaired bases through which the polynucleotide can take part in homotypic interactions to form oligomers.

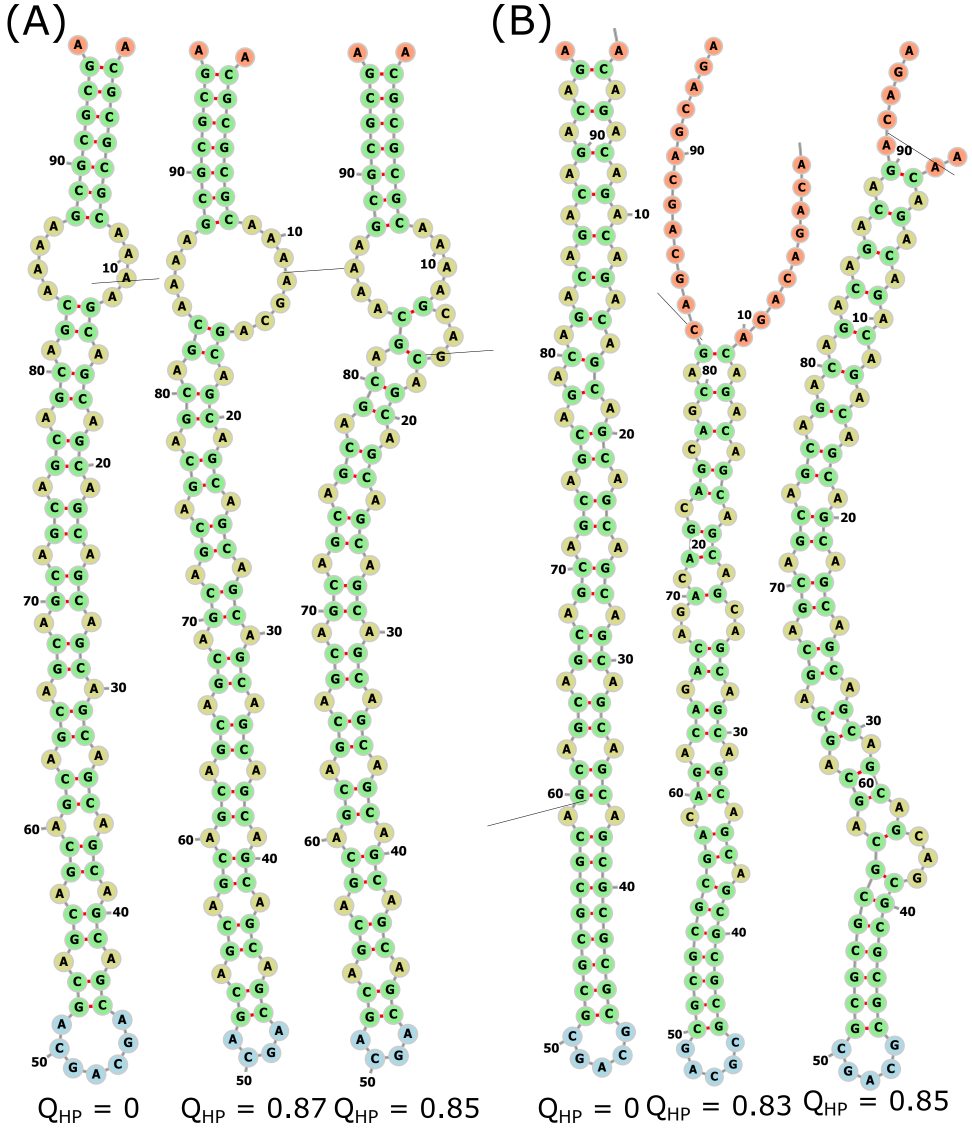


Figure S9: Conformations of scrambled sequences: (A) The GS and the excited states structures of the hairpins obtained for SS1. Corresponding $Q_{HP}$ values of the hairpins are denoted below. (B) Different hairpin structures for SS2. The GS with $Q_{HP} = 0$ contains no overhang at the terminal. The excited state, with $Q_{HP} = 0.83$ and 0.85, has overhangs through which the hairpin can interact with other RNA molecules to form higher order oligomers.

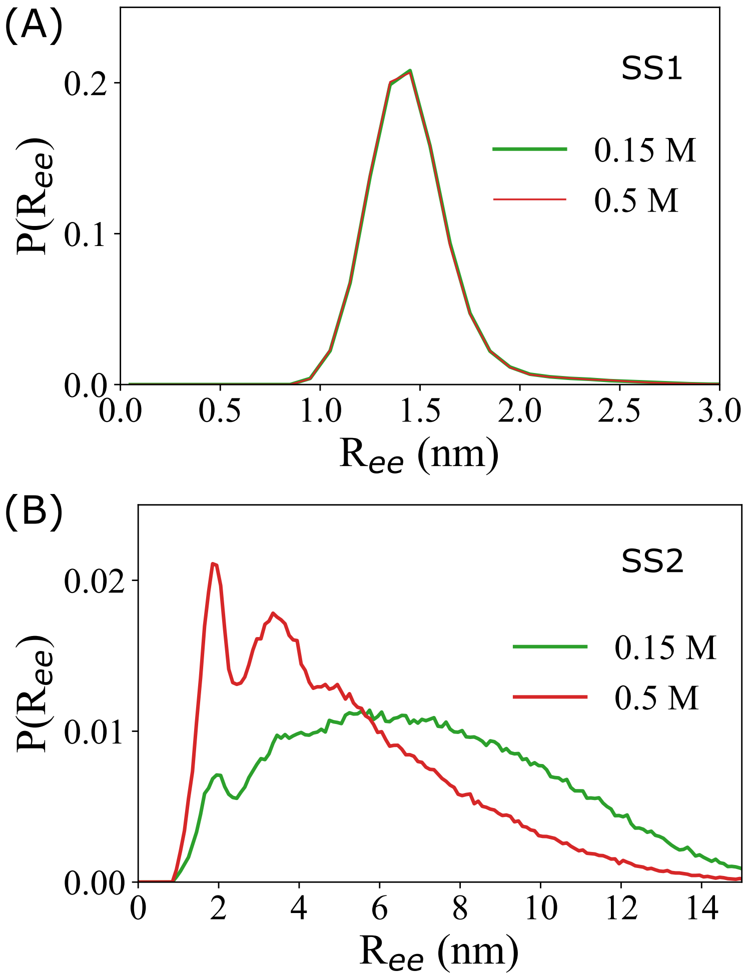


Figure S10: Distribution of end-to-end distances, $R_{ee}$, $P(R_{ee})$: (A) $P(R_{ee})$ at $C_S = 0.15$ and 0.5 M for SS1 are in green and red, respectively. The distribution has a peak with no slippage in the strand. (B) $P(R_{ee})$ for SS2 at $C_S = 0.15$ M and 0.5 M are in green and red, respectively. The distribution of $R_{ee}$ shifts to lower $R_{ee}$ value as the $C_S$ is increased, indicating slippage probability is reduced with an increase in $C_S$.
